# 3D modeling of CpG DNA binding with matrix lumican shows leucine-rich repeat motif involvement as in TLR9- CpG DNA interactions

**DOI:** 10.1101/2023.08.21.554201

**Authors:** Tansol Choi, George Maiti, Shukti Chakravarti

**Author notes:** Correspondence: GM,; SC.

## Abstract

Lumican is an extracellular matrix proteoglycan, known to regulate toll-like receptor (TLR) signaling in innate immune cells. In experimental settings, lumican suppresses TLR9 signaling by binding to, and sequestering its synthetic ligand, CpG-DNA, in non-signal permissive endosomes. However, the molecular details of lumican interactions with CpG-DNA are obscure. Here, the 3-D structure of the 22 base-long CpG-DNA (CpG ODN_2395) bound to lumican or TLR9 were modeled using homology modeling and docking methods. Some of the TLR9-CpG ODN_2395 features predicted by our model are consistent with the previously reported TLR9-CpG DNA crystal structure, substantiating our current analysis. Our modeling indicated a smaller buried surface area for lumican-CpG ODN_2395 (1803 A^2^) compared to that of TLR9-CpG ODN_2395 (2094 A^2^), implying a potentially lower binding strength for lumican and CpG-DNA than TLR9 and CpG-DNA. The docking analysis identified 32 amino acids in lumican LRR1-11 interacting with CpG ODN_2395, primarily through hydrogen bonding, salt-bridges and hydrophobic interactions. Our study provides molecular insights into lumican and CpG-DNA interactions that may lead to molecular targets for modulating TLR9 mediated inflammation and autoimmunity.

## 1. Introduction

Lumican is a member of the small leucine rich repeat proteoglycan (SLRP) family, widely present in various extracellular matrix (ECM) types (1, 2). A few SLRPs are post-translationally modified with glycosaminoglycan side chains in mature ECMs. Lumican is a modified keratan sulfate (KS) proteoglycan in connective tissues like the tendon, cornea and cartilage, but it is secreted as a simple glycoprotein by activated fibroblasts during infections and inflammation (3–5). Much of the core proteins consist of tandem repeats of leucine-rich repeat (LRR) motifs. A body of work, including transmission electron microscopy of collagen-rich tissues, recombinant core proteins and biochemical approaches show that lumican and other SLRP members associate with collagen fibrils to restrict their lateral growth and assembly, and that they localize to different bands on the fibrils (6–10). Functionally, the SLRPs regulate collagen fibril growth and proper assembly of ECMs, such that SLRP-null mice develop abnormal connective tissues (11–14). SLRP interactions with collagen fibrils engage multiple LRR repeats in the core proteins (15, 16). In addition to collagen-fibril regulations, emerging studies indicate that SLRPs interact with cell surface receptors, chemokines and pathogen associated molecular patterns, cementing their significant involvement with host immune responses (2, 4). Given that the SLRP core proteins are largely comprised of LRR motifs, they are members of the LRR superfamily of proteins which includes pathogen recognition toll-like receptors (TLRs).

Host pathogen recognition receptors (PRR) include TLRs that mediate host response to microbial pathogen associated molecular patterns, as well as certain host-derived danger associated molecular patterns (17, 18). Human and mice have 10 and 12 TLRs in humans and mice, respectively. TLRs 1, 2, 4, 5 localize to the plasma membrane, whereas TLR3, 7/8 and 9 are destined to endosomal compartments for PAMP recognition and signaling (18). TLR9 recognizes viral double stranded (ds) DNA in acidic endolysosomal compartments. TLR9 may also interact with host nuclear DNA as well as mitochondrial DNA released from dying cells during infections and inflammation, which contributes to increased risk for autoimmunity (19). Adequate recognition of pathogenic, but minimal recognition self-DNA, is achieved through guided trafficking and cleavage of TLR9 in endolysosomes. Our experimental study shows that ECM proteins like lumican may modulate recognition of pathogenic DNA (5). Future studies will also address whether ECM proteins minimize recognition of host-derived DNA by TLR9. Lumican and TLR9 have multiple tandem repeats of LRR motifs. LRR motifs from TLR9 have already been shown to interact with some synthetic forms of DNA ligands (20, 21). Our 3D modeling of lumican binding to DNA ligands, for the first time, identifies candidate DNA-binding LRR motifs in lumican.

LRR motifs, first discovered in 1985, are approximately 24 amino acid long highly conserved leucine rich stretches, with regularly spaced hydrophobic residues (22). Over 6000 proteins having LRRs are listed in PFAM, PRINTS, SMART, and InterPro databases (23). These LRR domains have a highly conserved segment (HCS) and a variable segment (VS) (24, 25) . The HCS, LxxLxLxxNxL, contains conserved leucine residues (Leu), which may be substituted with Isoleucine (lle), phenylalanine (Phe), or valine (Val). The ninth residue is usually an asparagine (Asn), or serine (Ser), threonine (Thr), or cysteine (Cys). LRR superfamily proteins generally have a horseshoe shape, where the Leu, Ile, Phe and Val at position 1, 4, 6, and 11 of HCS form the hydrophobic core of the protein comprised of beta sheets (24). The Asn at position 9 connects LRR to each other by forming hydrogen bond. The VS part forms loop and helix conformation at the outer convex surface (24).

In addition to the shared structural similarities discussed above, the SLRP core proteins harbor some TLR-like functions, and interact with pathogen recognition signals. In a recent study we found that macrophages from lumican-null mice show increased innate immune response to CpG DNA, a ligand for endosomal TLR9, while addition of recombinant lumican suppressed TLR9 response (5). The study further demonstrated that exogenous lumican and CpG DNA co-localized in early endosomes, which are not enriched for TLR9. In *in vitro* recombinant lumican binds CpG DNA in a dose-dependent manner. The data collectively indicated that lumican binds CpG DNA and sequesters it in TLR9-poor endosomes to restrict response to DNA. However, the molecular underpinnings in the lumican-DNA interactions are unknown.

Here, we used *in-silico* approaches to predict the LRR sequences in lumican and their mode of interactions with CpG DNA. We modeled lumican-CpG DNA interactions using the 22-mer synthetic CpG DNA, henceforth termed as CpG ODN_2395 that has been shown to experimentally stimulate mouse and human TLR9 responses. A few in silico studies have examined TLR9 interactions with CpG DNA forms other than the CpG ODN_2395 we studied here (21, 26). Prior crystallographic studies on TLR9 extracellular domain (TLR9 ECD) used a variation of the 12-mer synthetic CpG ODN_1668, CATGACGTTCCT and a 6-mer that binds to a second DNA binding region to enhance signals in human, equine and bovine TLR9 (27). However, there are no experimental or model predictions of TLR9 interactions with the 22-mer CpG ODN_2395 ligand. Therefore, to ascertain the extent to which lumican mimics TLR9 in its interactions with the CpG ODN_2395 ligand, we also modeled mouse TLR9 ECD interactions with this ligand. Our studies provide new insights into matrix lumican and mouse TLR9 interactions with a CpG ODN_2395 known to evoke strong innate immune response *in vivo*.

## 2. Results

### 2.1. 3D modeling of lumican and LRR motif-carrying domains in Lumican and TLR9

Because mouse lumican has not been crystallized, we used the AlphaFold Protein Structure Data Base (37, 38) to visualize the predicted structure of mouse lumican in PDB format. We also superimposed TLR9 LRR1-13 on the lumican model for visual comparison (**Fig. 1A**). The analysis range of lumican (amino acid residues 40-309) yielded a high per residue confidence metric or predicted local distance difference test (pLDDT) score of more than 90%, and a predicted aligned error (PRE) ranging from 0 to 15 A.

**Figure 1.**
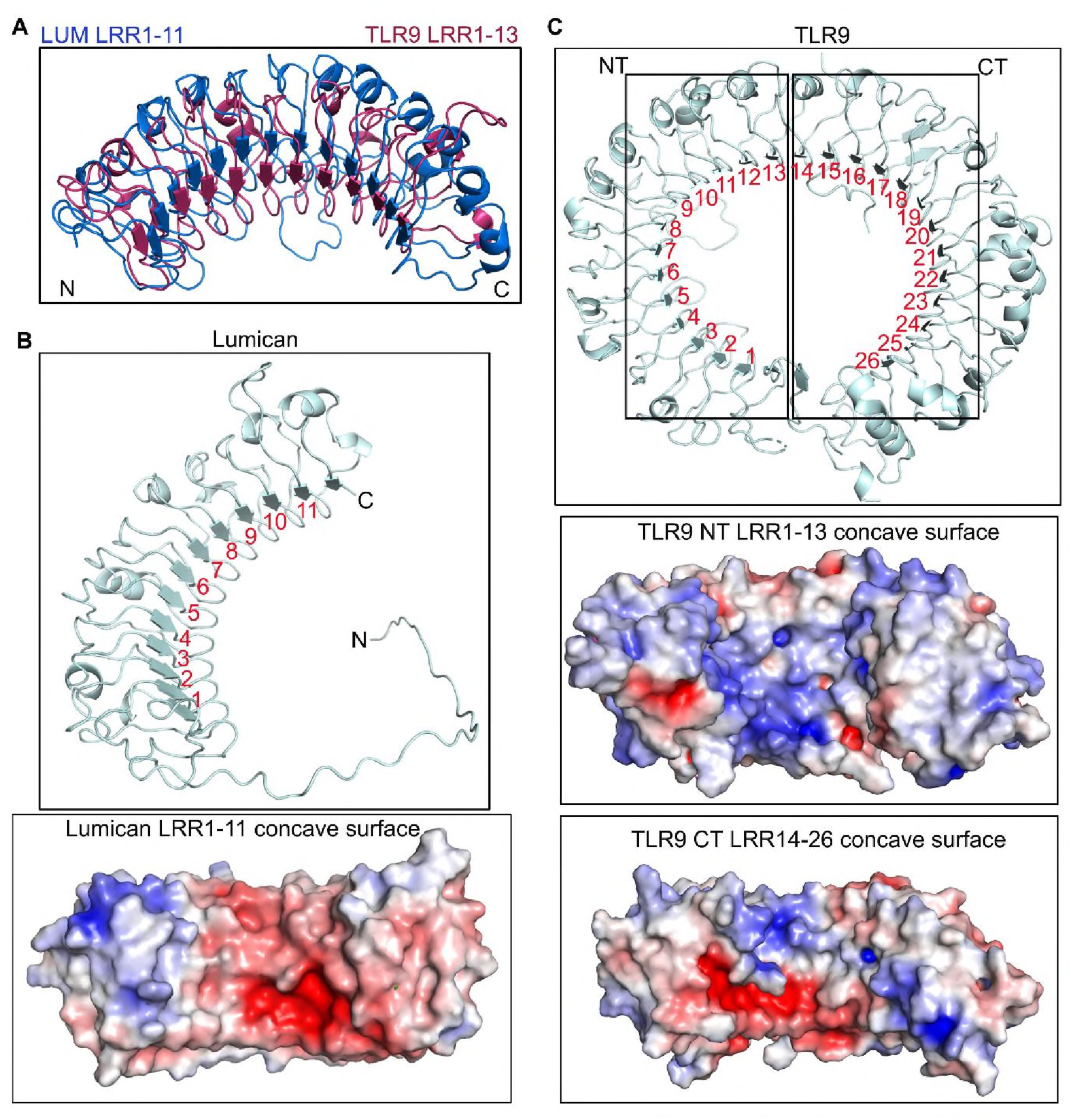
Lumican model structure and similarities with TLR9. **A.** The model depicts the structural alignment of TLR9 LRR1-13 and lumican LRR1-11 based on ClustalW pairwise sequence alignment. **B-C.** Horseshoe-like structure model and surface charge representation showing basic (blue) and acidic (red) properties of lumican concave surface showing LRR1-11 repeats **(B)** and TLR9 showing LRR1-26 repeats **(C)**.

The LRR motifs, based on their consensus sequence, can be categorized into seven classes bacterial (S), cysteine containing (CC), plant specific (PS), SDS22, ribonuclease inhibitor like (RI), irregular (I) and typical (T) (25). In a pairwise alignment using Clustal Omega, lumican shared an overall identity of 22% and a similarity of ∼40% with TLR9 (**Fig. S1)**. We detected the highest similarity between lumican LRR motifs 1-11 and LRR 1-13 of mouse TLR9 (**Fig. 1A)**. Lumican follows a typical pattern of LRR repeat consensus with approximately 20 residues per repeat, where the central ten LRR motifs (LRR1-11) are a mixture of T, S, SDS22 and PS classes, flanked by an N-terminal LRRNT motif with 4 cysteine residues and a C-terminal LRRCT motif with 2 cysteine residues **(Fig. S2)**. TLR9 has a total of 26 LRR motifs, where several are irregular, with insertions of variable lengths of amino acid that disrupt the average 26 residues per LRR structure. Thus, LRR 2, 5, 8, and 11 of mouse TLR9 have 10-16 insertions at position 10 after the 9^th^ consensus Asn residue (28). Surface charge distribution analysis along the concave surface of lumican shows that it consists of basic and acidic patches, and the involves equal number of basic and acidic residues acidic and basic interface residues **(Fig. 1B, Fig. S3A).** On the other hand, in TLR9, LRR1-13 and LRR13-26 motifs along the concave surface are predominantly basic patches **(Fig. 1C, Fig. S3B-C).**

### 2.2. CpG ODN_2395 docking and interaction surfaces on Lumican and TLR9 ECD

As TLR9 interactions with CpG ODN_2395 has not been studied before, we evaluated the docking of CpG ODN_2395 on the “unliganded form” (PDB: 3WPF) of mouse TLR9. Our docking analysis of CpG ODN_2395 on lumican identified LRR 1-11 (**Table.1**) and on LRR2-8, 13, 15, and 20-25 on TLR9-ECD **(Table.2)** as the most likely sites of interactions.

**Table 1.** Lumican -CpG ODN_2395 interacting amino acid residues.

| **Lum Interface** | **BSA** | **LRR#** | **LRR type** |
| --- | --- | --- | --- |
| SER45 | 17% | NT | - |
| ARG73 | 22% | LRR1 | T |
| ASN74 | 23% |  |  |
| ILE95 | 34% | LRR2 | T |
| ASP97 | 90% |  |  |
| HIS98 | 42% |  |  |
| LYS119 | 19% | LRR3 | PS |
| HIS121 | 65% |  |  |
| ASN123 | 72% |  |  |
| TYR124 | 20% |  |  |
| ASP140 | 59% | LRR4 | SDS22 |
| GLN142 | 96% |  |  |
| PHE162 | 42% | LRR5 | T |
| TYR164 | 99% |  |  |
| GLN166 | 67% |  |  |
| TYR187 | 94% | LRR6 | S |
| ASP189 | 58% |  |  |
| THE208 | 71% | LRR7 | T |
| TYR210 | 93% |  |  |
| ASP212 | 7% |  |  |
| TYR232 | 59% | LRR8 | SDS22 |
| ARG234 | 71% |  |  |
| SER236 | 84% |  |  |
| HIS237 | 15% |  |  |
| GLU257 | 77% | LRR9 | SDS22 |
| ASP259 | 94% |  |  |
| SER261 | 100% |  |  |
| TYR262 | 41% |  |  |
| TYR279 | 86% | LRR10 | Irregular |
| GLU281 | 94% |  |  |
| VAL282 | 2% |  |  |
| ARG309 | 37% | LRR11 | Irregular |

**Table 2.** TLR9 - CpG ODN_2395 interacting amino acid residues.

| **TLR9 Interface** | **BSA** | **LRR#** | **LRR type** |
| --- | --- | --- | --- |
| LYS95 | 29% | LRR2 | Irregular |
| ASN129 | 57% | LRR3 | S |
| SER131 | 100% |  |  |
| TYR132 | 57% |  |  |
| SER149 | 66% | - | - |
| SER151 | 85% | LRR4 | T |
| HIS152 | 70% |  |  |
| VAL171 | 48% | LRR5 | Irregular |
| PHE173 | 79% |  |  |
| ASP175 | 84% |  |  |
| TYR179 | 63% |  |  |
| TYR180 | 23% |  |  |
| LYS181 | 9% |  |  |
| HIS203 | 66% | LRR6 | S |
| SER205 | 0% |  |  |
| LYS207 | 67% |  |  |
| TYR208 | 56% |  |  |
| TYR224 | 52% | LRR7 | T |
| LEU226 | 48% |  |  |
| PRO262 | 17% | LRR8 | Irregular |
| ASN263 | 48% |  |  |
| GLN399 | 55% | LRR13 | T |
| MET400 | 47% |  |  |
| PHE467 | 37% | - |  |
| ARG482 | 24% | LRR15 | T |
| ARG613 | 36% | LRR20 | Irregular |
| ASN640 | 52% | LRR21 | T |
| HIS642 | 51% |  |  |
| ARG662 | 23% | LRR22 | T |
| TYR665 | 50% |  |  |
| GLN689 | 15% | LRR23 | SDS22 |
| SER711 | 14% | LRR24 | T |
| HIS735 | 20% | LRR25 | T |

We next visualized CpG ODN_2395 interactions with lumican and TLR9 using 3D modeling. A large area along the concave surface of lumican engages with CpG ODN_2395, where LRR2-8 from the N-terminal end forms a binding groove for the 5’ end of CpG ODN_2395 **(Fig. 2A-B)**.

**Figure 2.**
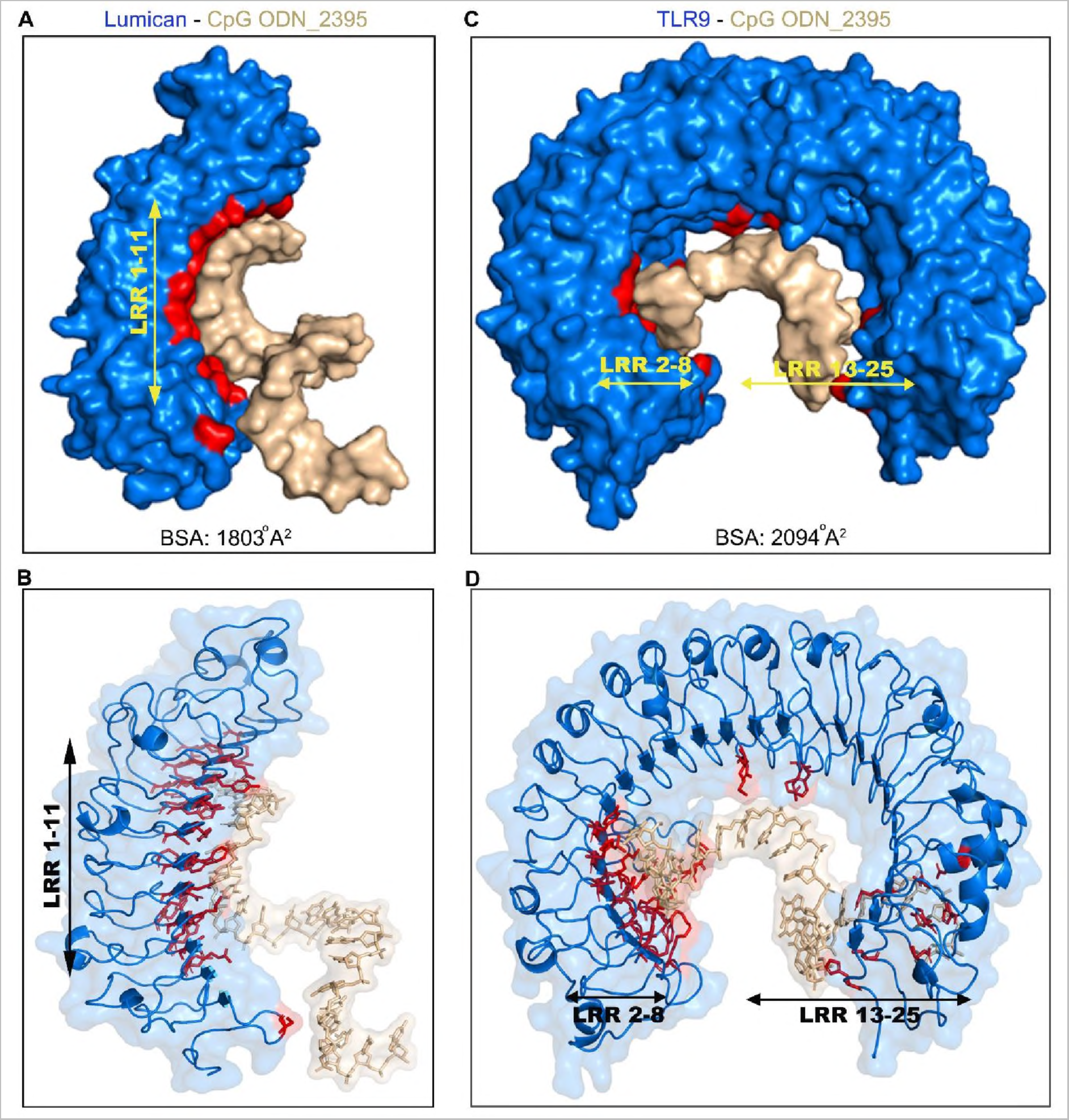
Modeling of CpG ODN_2395 bound Lumican. **A.** Surface view of lumican-CpG ODN_2395 complex shows the buried surface area (BSA) in red, lumican (blue), CpG ODN_2395 (golden). The combined BSA for Lumican-CpG DNA complex is 1803 A^2^. Lumican residues contribute 42% (761 A^2^) of the BSA whereas CpG OdN_2395 contributes 58% (1042 A^2^). **B.** A ribbon 3D model showing all lumican residues (red) that interact with CpG ODN_2395 (golden). **C.** Surface view of the TLR9-CpG DNA complex shows the buried surface area (BsA) in red, TLR9 (blue) and CpG ODN_2395 (golden). The combined BSA for TLR9-CpG DNA complex is 2094 A^2^. TLR9 residues contribute 42% (878 A^2^) of BSA. **D.** A ribbon 3D model showing all TLR9 residues (red) that interact with the CpG ODN_2395 (golden).

We also visualized the CpG ODN_2395 interacting surfaces in TLR9 ECD. The binding region includes LRR2-13 from the N-terminus and LRR15, 20-25 from the C-terminus **(Fig. 2C-D, Table.2).** This is different from those reported earlier for interactions of TLR9 ECD with the shorter CpG ODN_1668 where only LRR1-8 were engaged from the N-terminus and LRR20-22 from C-terminus **(Fig. S4)**. We further examined the predicted buried surface area (BSA), defined as the surface area buried or unavailable to solvents in protein bound to CpG DNA. Analysis of the lumican - CpG ODN_2395 interaction yielded a BSA of 1803 A^2^ **(Fig. 2A)**; whereas the TLR9 ECD-CpG ODN_2395 interaction yielded a BSA of 2094 A^2^ **(Fig. 2B)**. This indicates that compared to lumican, a slightly larger surface area of TLR9 engages with CpG ODN_2395. By contrast, a previous study on TLR9 ECD interactions with the smaller 12 mer CpG ODN_1668 showed 1430 A^2^ BSA, predictably suggesting smaller surface occupancy by the smaller CpG DNA (21).

### 2.3. Lumican and CpG ODN_2395 interactions involve hydrogen bonding, salt-bridges and hydrophobic interactions

The lumican-CpG ODN_2395 interface involves 32 amino acid residues from lumican, and 9 nucleotide bases from the CpG ODN_2395, at an average resolution of 3.3 A. The interacting atoms from the bases in CpG ODN_2395 and the amino acids from lumican are shown **(Fig. 3A, Table.3)**. The interacting residues include 7 basic (Lys119, His98, His121, His237, Arg73, Arg234, Arg309) residues and two hydrophilic asparagine residues, Asn74, Asn123, seven acidic residues (Glu257, Glu281, Asp97, Asp140, Asp189, Asp212, Asp259) and 16 other hydrophobic and polar uncharged amino acid residues. The interactions include 11 hydrogen bonding followed by 4 salt-bridge and 2 hydrophobic interactions **(Fig. 3A, Table.3)**. Hydrogen bonding interactions with CpG ODN_2395 pyrimidine/purine bases are the predominant, where 82% H-bond donor residues reside in lumican, and 82% of H-bond acceptors reside in CpG ODN_2395 **(Table.3).** Biochemically, the amino acids in lumican that form the 11 H-bonds include Asn74, Arg234 and Arg309, two acidic Asp259 and Glu281,a neutral Ser261, two acidic Gln142, Gln166, and three aromatic Tyr164, Tyr187 and Tyr210. The four salt bridges are formed between opposite charges in polar/charged residues including Glu257, Asp189, His98 and His121. Acidic residues Glu257 and Asp189 form salt bridges with the G3 and G6 purine bases of CpG ODN_2395, respectively **(Fig. 3B-D)**. However, the positive charge centers of the bases form electrostatic interactions with the Glu257 and Asp189 negative charges of the carboxyl side chains **(Fig. 3D)**. The positive imidazole side chain of His98 and 121 forms salt-bridges with the ribose-phosphate backbones of CpG ODN_2395. Hydrophobic/pi-stacking interactions are observed between aromatic ring of Tyr 262 and the pyrimidine base ring of T1, where the Tyr aromatic ring is perpendicular to thymidine base ring making it ideal for pi-stacking interactions **(Fig. 3C).**

**Figure 3.**
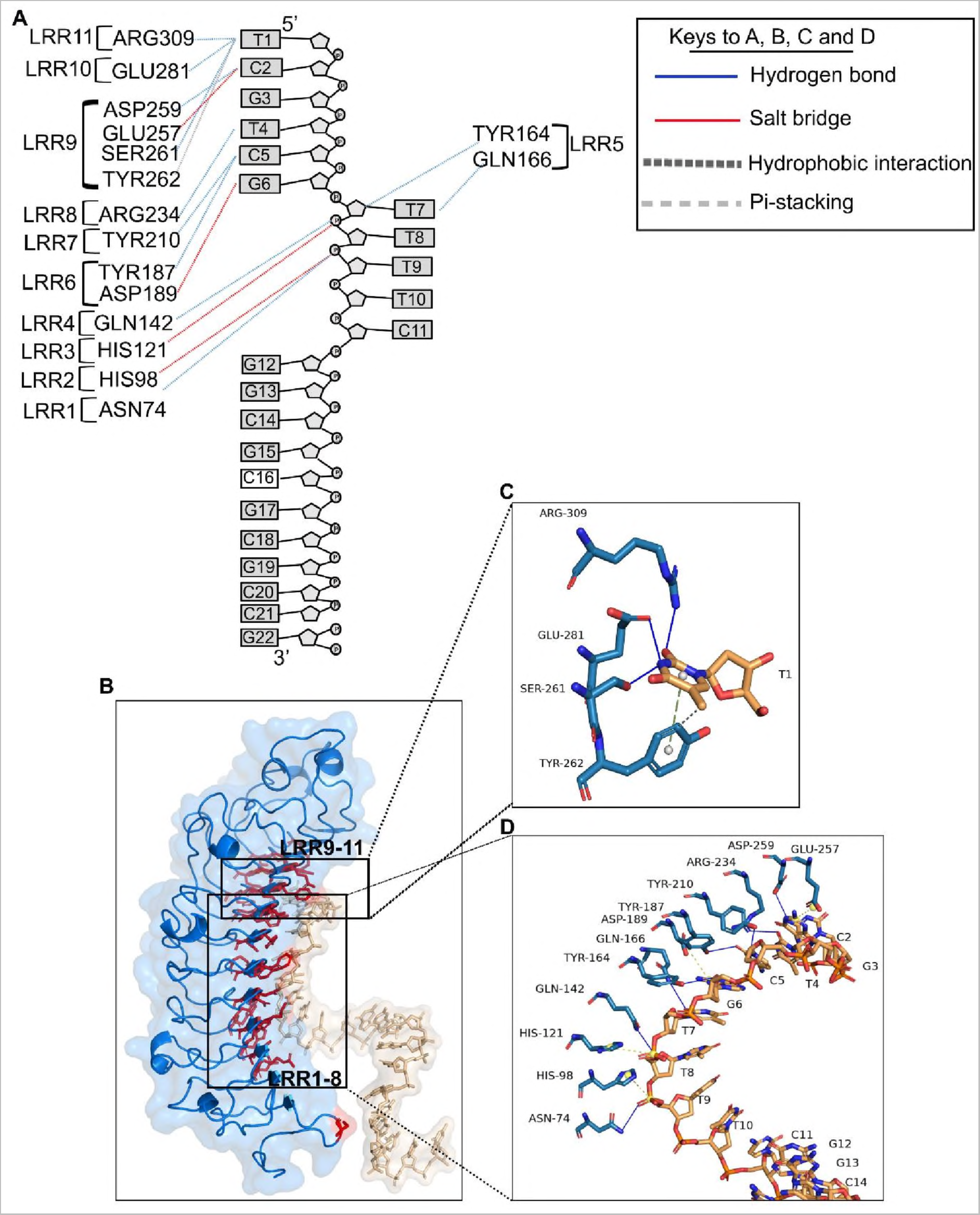
Close up view Lumican and CpG ODN_2395 interactions. **A.** A schematic diagram showing lumican interacting LRRs from lumican and nucleotides from CpG ODN_2395. The dotted lines in blue indicate H-bonds, red/yellow salt-bridges and black pi-stacking interactions. **B.** A 3D ribbon model showing all lumican residues (red) that interact with the CpG ODN_2395 (golden). **C-D.** A stick model showing interaction of lumican with the first 5’ thymidine (T) base **(C)** and the next 8 bases of CpG ODN_2395 **(D)**. The yellow ball shows the charge centers in CpG ODN_2395 and the grey ball indicates the aromatic charge centers of amino acids.

**Table 3.** Different types of interaction between lumican and CpG ODN_2395.

| Lum-CpG |  |  |  |  |
| --- | --- | --- | --- | --- |
| Hydrogen bond | CpG ODN_2395 | Atom | Lum | Atom |
|  | T1 | N3 | GLU281 | OE1 |
|  |  | N3 | SER261 | OG |
|  |  | O2 | ARG309 | NH2 |
|  | C2 | N4 | ASP259 | OD2 |
|  | T4 | O2 | ARG234 | NH1 |
|  | C5 | N3 | TYR210 | OH |
|  |  | O2 | TYR187 | OH |
|  | G6 | O4 | TYR164 | OH |
|  | T7 | O2 | GLN166 | NE2 |
|  | T8 | O5 | GLN142 | NE2 |
|  | T9 | OP1 | ASN74 | ND2 |
| Salt bridge | CpG ODN_2395 | Charge center | Lum | charge center |
|  | G3 | Purine ring | GLU257 | carboxylic |
|  | G6 | Purine ring | ASP189 | carboxylic |
|  | T8 | P-backbone | HIS121 | imidazole |
|  | T9 | P-backbone | HIS98 | imidazole |
| Van der Waals | CpG ODN_2395 | Atom | Lum | Atom |
|  | T1 | C7 | TYR262 | CE2 |
| Pi-stacking | CpG ODN_2395 | Atom | Lum | Atom |
|  | T1 | Pyrimidine ring | TYR262 | Tyrosyl ring |

H-BOND DONOR
H-BOND ACCEPTOR

The hydrophobic and the basic residues of lumican form the interface with the first and third groove of CpG ODN_2395. The lumican residues that forms close contact with the first grove of CpG ODN_2395 includes Asp189, Asp259, Glu257, Glu28, Arg234, Arg309 and Tyr164, Tyr187, Tyr210 and Ser261 that forms salt bridges/H-bonds with the two 5’ CpG repeats (5’-TCGTCG). These “hot spot” lumican residues in LRR5-11 forms a groove that interacts with the two 5’ CpG repeats. These lumican residues, therefore, are biochemically somewhat similar to the TLR9 ECD LRR2-13, as predicted by the pairwise alignment score **(Fig. S5A-C).**

### 2.4. TLR9 ECD interactions with CpG ODN_2395 involve N-terminal LRR2—13 motifs

The TLR9-CpG ODN_2395 interface encompasses 33 residues of TLR9 and 10 nucleic acids of CpG ODN_2395 at an average resolution of 3.5 A **(Table.2, Fig. 2C-D).** As indicated by the APBS map, the interface residues involve many basic residues that form distinct basic patches in N and C terminal side of TLR9 (**Fig. S3B-C)**. The N-terminal LRR2-13 of TLR9 is contributing the most towards the binding affinity, comprising 553 Å^2^ out of 878 Å^2^ of the buried surface area of TLR9 (**Fig. 2C**). The TLR9 N-terminal LRR2-13 region contains seven basic residues Lys95, Asn129, His152, Lys181, His203, Lys207 and Asn263, and one acidic residue Asp175 **(Fig. S3B)**. There are 15 additional neutral, polar or hydrophobic (Ser131, Ser149, Ser151, Ser205, Tyr132, Tyr179-180, Tyr208, Tyr224, Val171, Phe173, Leu226, Pro262, Gln399 and Met400) residues in these interacting LRRs (**Table.2**). However, the C-terminal LRR15-25 contains six basic residues Arg482, Arg613, Asn640, His642, Arg662 and His735 **(Fig. S3C)**. Our results show that the predominant type of interaction for TLR9-CpG ODN_2395 is hydrogen bonding, involving 12 H-bond interactions where 75% of the H-bond donors reside in TLR9 and 75% of the acceptors in CpG ODN_2395. Specifically, the H-bond donors are four tyrosine residues (Tyr132, Tyr179, Tyr180 and Tyr665) and other residues (His203, Arg613, Asn263, Ser131, Ser205 and Lys207) (**Fig. 4A-E**). The salt bridge interactions are the next prevailing interaction, mostly involving charge-charge interactions between positively charged basic residues (Lys95, His152, Arg482, Arg613, His642 and His735) and negatively charged sugar-phosphate backbone atoms **(Fig.4A, Table.4)**. One exception was the acidic residue Asp175, where the carboxylic side chain forms a salt-bridge with the positive charge center of the G3 purine base ring (**Fig. 4B-C**). Overall, 11 out of the 18 bonding interactions including H-bond/salt-bridges are from LRR2-13. The N-terminus LRR2-13 of the TLR9-CpG ODN_2395 interface comprises 63% of the BSA (878 A^2^), while LRR15-26 from C-terminus makes up only 37% of BSA (878 A^2^) (**Table.2**). Thus, the C-terminus of LRR15-26 has low binding affinity for CpG ODN_2395, which is consistent with experimental data that shows its reduced involvement in TLR9 downstream signals (20). The His152 and His203, and Tyr132, Tyr179 and Tyr180, Ser131 and Ser205, and Lys207 and Asn263 of LRR2-8 interacts closely with the 5’ end hexamer (5’-TCGTCG) of CpG ODN_2395 forming the H-bonds and salt bridges **(Fig. 4B-C)**. Additional interacting atoms from CpG ODN_2395 are also shown (**Fig. 4D-E, Table.4**)

**Figure 4.**
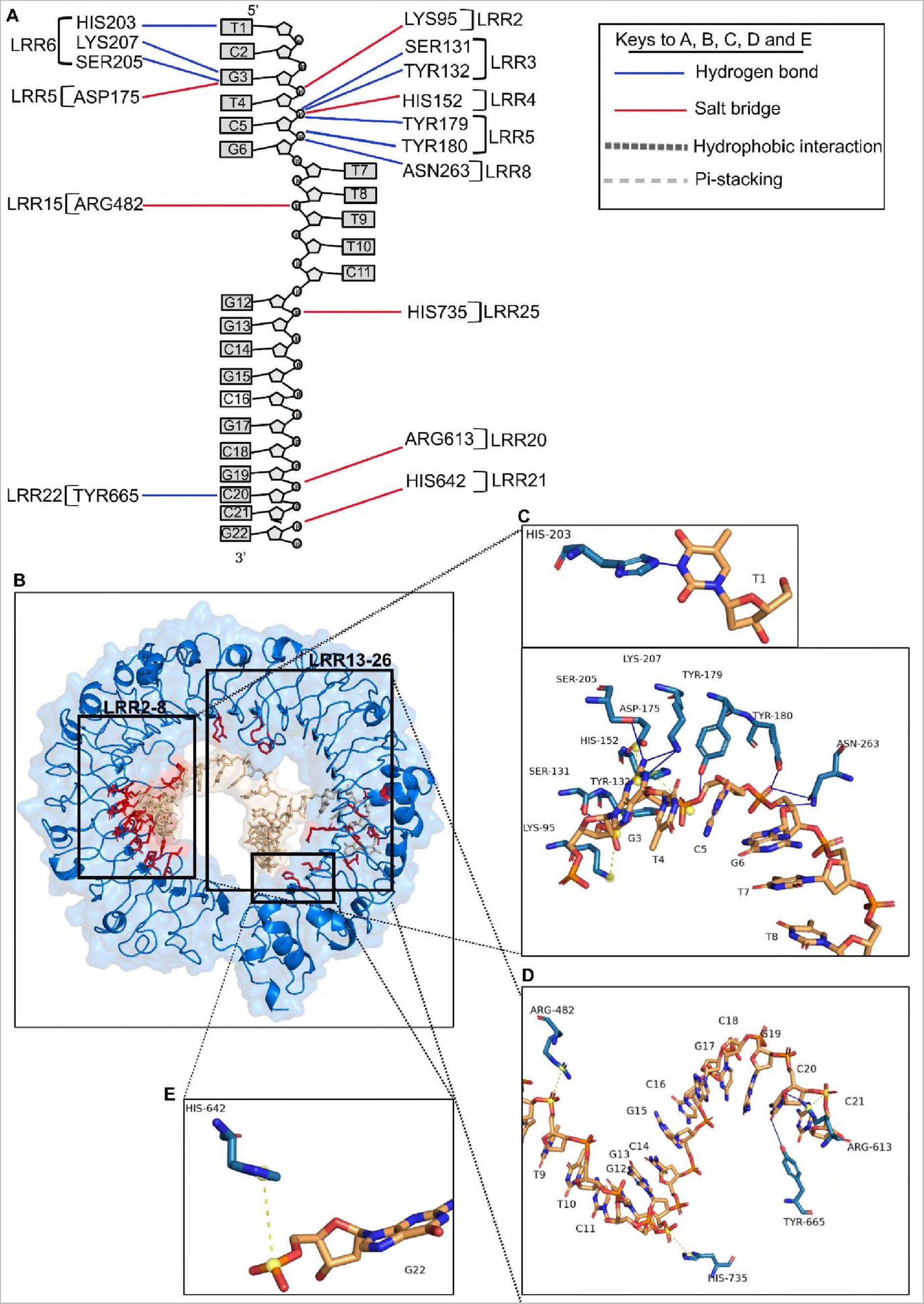
Close up view of TLR9 ECD and CpG ODN_2395 interactions. **A** A schematic diagram showing all TLR9 LRR and CpG ODN_2395 interactions. The dotted lines in blue indicates H-bonds and red/yellow salt-bridges. **B.** A 3D ribbon model showing all the TLR9 residues (red) that interacts with the CpG ODN_2395 (golden). **C-E.** Stick model indicates the interaction of TLR9 LRR2-8 with the bases or sugar phosphate backbone of the 5’-TCGTCG of CpG ODN_2395 **(C)**, the LRR15, 20-22 and 25 interacting with phosphate backbone of the thymidine (T) at position 8, guanosine (G) at position 13 and cytosine (C) at position 20 of CpG ODN_2395, respectively **(D)** and LRR21 interacting with the phosphate backbone of guanosine (G) at position 22 of CpG ODN_2395 **(E)**. The yellow ball shows the charge centers in CpG ODN_2395.

**Table 4.** Different types of interaction between TLR9 and CpG ODN_2395.

| TLR9-CpG |  |  |  |  |
| --- | --- | --- | --- | --- |
| Hydrogen bond | CpG ODN_2395 | Atom | TLR9 | Atom |
|  | T1 | N3 | HIS203 | NE2 |
|  | G3 | N1 | LYS207 | NZ |
|  |  | N2 |  | NZ |
|  |  | N2 | SER205 | OG |
|  | T4 | OP1 | SER131 | OG |
|  |  | O3 | TYR132 | OH |
|  | C5 | O5 | TYR179 | OH |
|  | G6 | OP1 | TYR180 | OH |
|  |  | OP1 | ASN263 | OD1 |
|  |  | O5 |  | ND2 |
|  | C20 | O2 | TYR665 | OH |
|  |  | O4 | ARG613 | NH2 |
| Salt bridge | CpG ODN_2395 | Charge center | TLR9 | Charge center |
|  | G3 | Purine ring | ASP175 | carboxylic |
|  | T4 | P-backbone | LYS95 | amine |
|  | C5 | P-backbone | HIS152 | imidazole |
|  | T9 | P-backbone | ARG482 | amine |
|  | G13 | P-backbone | HIS735 | imidazole |
|  | C21 | P-backbone | ARG613 | amine |
|  | G22 | P-backbone | HIS642 | Imidazole |

H-BOND DONOR
H-BOND ACCEPTOR

As indicated by previously, TLR9 interacting residues with CpG ODN_1668 12mer are clustered in LRR1-8 from the N-terminus, which includes the generic PAMP binding surface LRR2, 5, and 8, followed by LRR20-22 from the C-terminus (21). The TLR9-CpG ODN_1668 12mer interaction region include N-terminus basic patches of TLR9 that engage in hydrogen bonding and satisfies charge-complementarity to negatively charged ribose-phosphate backbone **(Fig. S4)**. The prevailing interactions between LRR1-8 and ODN_1668 are H-bonding between sugar-phosphate backbones.

### 2.5. The predicted effects of collagen type I on lumican interactions with CpG ODN_2395

Multiple studies have shown that lumican associations with collagen type I regulate collagen fibrillogenesis *in vitro* (7), and that similar interactions *in vivo* impact collagen fibril assembly (12, 15, 29). One study found that synthetic peptides from the lumican LRR5-7 region competes with full length lumican and significantly compromises collagen type 1 fibrillogenesis (9). Moreover, the D212N recombinant lumican which carries this mutation in LRR5-7, disrupts both binding of lumican to collagen and subsequent collagen fibrillogenesis (9). Our simulation shows that lumican upon binding to CpG ODN_2395 creates significant amount of BSA (760 A^2^ of 1803 A^2^, which is 42%). The BSA of most bona-fide protein-ligand interactions are observed within this range (30). Our prior interface analysis identified Asp212, Gln166, Tyr210, Asp189, Tyr164, Thr208, Tyr187 and Phe162 as interacting with CpG ODN_2395, and these fall within the LRR5-7 that interact with collagen **(Fig. 5A)**. The residue Asp212 (D212) within LRR7 which is involved in fibrillogenesis is predicted to be buried upon CpG binding. This site is also highly conserved in lumican from different species as shown by multiple sequence alignment, indicating its functional importance **(Fig. 5B)**. These analyses also imply that collagen binding will likely interfere with the ability of LRR5-7 to bind CpG-DNA.

**Figure 5.**
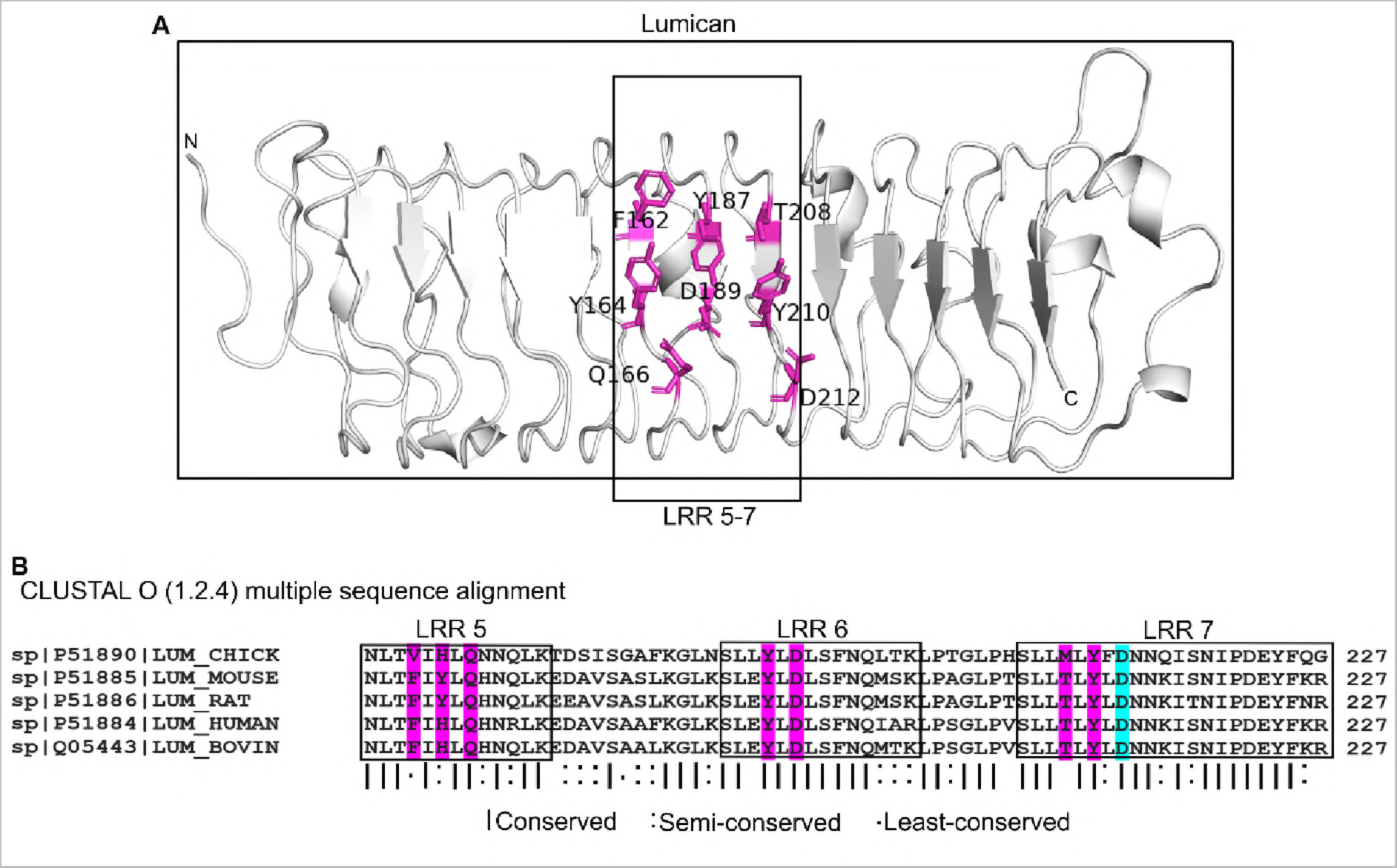
CpG DNA binding sites in LRR5-7 of lumican also known to bind collagen I. **A.** CpG ODN_2395 binding residues in LRR5-7 of lumican. CpG ODN_2395 binding residues colored magenta. **B.** Multiple sequence alignment of lumican LRR5-7 from different species. Interface residues are marked in magenta. Asp212 (D212) in cyan is the interface residue that interacts with collagen type I

## 3. Discussion

Our earlier experimental studies showed that lumican binds CpG ODN_2395, a synthetic ligand for TLR9 to restrict response and subsequent induction of proinflammatory cytokines by TLR9 (5). Here, for the first time, we used *in silico* approaches to identify the DNA-interacting region within the core protein of lumican at a molecular level, and characterize the types of bonding involved in these interactions. Like other LRR family members, the lumican core protein is made up almost entirely of tandem repeats of LRR motifs. Our analysis shows that while overall identity (22%) and similarity (∼40%) between mouse lumican and TLR9 were not high, the lumican LRR motifs, except its last LRRCE motif share the strongest similarity with the anterior half of TLR9, namely, LRR 1-13. Like other LRR proteins, lumican and TLR9 have a solenoid shape, where highly conserved residues are mainly located in the concave surfaces. Interactions with CpG DNA largely occur through the concave surface of the solenoid structures. Both proteins have a mixture of T, S, SDS22 and PS classes of LRRs, where, S and T are considered to interact with proteins and DNA and have role in immunity and defense. Indeed, lumican LRRs 1, 2 and 5-7, which are mixture of S and T class, are predicted to engage with CpG ODN_2395.

Analysis of BSA in the interactions of lumican (1803 A^2^) or TLR9 ECD (2094 A^2^) with CpG ODN_2395 suggest that, compared to TLR9 ECD, a slightly lower surface area of lumican is involved. This may imply a weaker binding strength between lumican and CpG ODN_2395 than between TLR9 ECD and CpG ODN_2395. However, from a functional standpoint in tissues, ECM proteins like lumican are present at a much greater abundance than pathogen recognition receptors, like TLR9, and thus expected to be impactful. Multiple, still incompletely understood mechanisms are in place to control TLR9 response to pathogenic DNA as well as self-DNA for controlled inflammatory responses to infections, and minimize harmful response to self-DNA (18, 19, 31–34). Synthesized in the endoplasmic reticulum, TLR9 is bound by Unc93B1, a membrane adaptor protein, which facilitates proper folding and incorporation into transport vesicles that apparently takes it to the plasma membrane first, before its final location in acidic endolysosomes where it is proteolytically cleaved to release a smaller functional protein (35). Our prior studies show that there is an abundance of lumican at the cell surface, and it is also endocytosed by immune cells (5). Therefore, binding of DNA ligands by lumican at the cell surface can counteract potential ligand interactions with TLR9 transiently located at the cell surface. We also showed that endocytosed lumican colocalizes with CpG DNA, and these endosomal sites are low in TLR9. This is another way that ligand interactions with TLR9 is reduced by lumican. We still do not know how lumican favors the separation of the DNA ligand from TLR9 in endosomes. The overall effects of lumican, and likely some of the other SLRPs, are to reduce ligand availability and interactions with TLR9 and to reduce inflammatory responses to pathogenic and self-DNA.

A potential limitation of our study is that we used the lumican core protein, instead of the glycoprotein or the proteoglycan forms of lumican in our 3D modeling. The negatively charged KS side chains of lumican will impact interactions of the core protein. However, we have shown experimentally, that during infections and inflammation, much of the lumican interacting with macrophages is not the proteoglycan, but the simple glycoprotein form without the KS side chains.

In summary, our study provides molecular details in the interactions between lumican and a synthetic ligand for TLR9, providing an avenue for future development of drug targets. Additional studies are required to expand our knowledge of the full extent of natural TLR9 ligands that lumican and other SLRPs interact with to regulate inflammation and autoimmunity.

## 4. Materials and Methods

### 4.1. Data sources

FASTA sequences of mouse TLR9 and lumican were obtained from UniProt with primary accession number Q9EQU3 and P51885, respectively. The Class C CpG ODN_2395 sequence (5’-TCGTCGTTTTCGGCGCGCGCCG-3’) was obtained from Invivogen (#tlrl-2395). We used the crystal structure of mouse TLR9 extracellular domain (TLR9 ECD) not bound to a ligand “unliganded form” (PDB:3WPF) obtained from Protein Data Bank (wwPDB) (36). Because mouse lumican has not been crystallized, we used the AlphaFold Protein Structure Data Base (37, 38) to visualize the predicted structure of mouse lumican in PDB format. The analysis range of lumican (amino acid residues 40-309) yielded a high pLDDT score of more than 90%, and a predicted aligned error (PRE) ranging from 0 to 15 A.

### 4.2. TLR9 and lumican local similarity analysis

Lumican and TLR9 FASTA sequences from UniProt were submitted to Clustal Omega local alignment tools (39) to align the most similar regions between two sequences. The lumican sequence 3-321 corresponding LRR1-11 out of 338 and TLR9 sequence 19-408 corresponding LRR1-13 out of 1032 were aligned. We introduced 121 gaps to align both sequences with default gap opening penalty score 10 and extension penalty score 0.5. The matched sequences have 165 conserved and semi-conserved residues out of 415, showing 22% sequence identity and 40% sequence similarity with alignment score 225.5 after subtracting gap opening and extension penalty score.

The results are the following:

- Aligned_sequences: LUM_MOUSE and TLR9_MOUSE
- Gap_penalty: 10.0
- Extend_penalty: 0.5
- Length: 415
- Identity: 92/415 (22.2%)
- Similarity: 165/415 (39.8%)
- Gaps: 121/415 (29.2%)
- Alignment Score: 225.5

The local structural alignment between two proteins were conducted using PyMOL Align command line (https://pymolwiki.org/index.php/Align) (40). It performs a local sequence alignment between two proteins using Clustal Omega local sequence alignment tool followed by structural superimposition with the cycles of refinement in order to reject structural outliers. The lumican LRR1-11 and TLR9 LRR1-13 totaling 1494 atoms were aligned with 5 cycles of structural refinement with the average distance between the atoms of two superimposed proteins or RMSD value 6.4. Structural outliers with the average RMSD= 6.83 were rejected during the cycles (53 atoms).

The following parameters were used to run PyMOL structural alignment between Lumican and TLR9:

- outlier rejection cycles: 5
- RMS outlier rejection cutoff: 2.0
- Mobile_state: 0=all states
- Target_state: 0=all states
- Transform: 1=superposition

The results obtained are following:

- Alignment score: 225.5
- ExecutiveAlign: 1819 atoms aligned.
- ExecutiveRMS: 53 atoms rejected during cycle 5 (RMSD=6.83)
- Executive: RMSD = 6.411 (1494 to 1494 atoms)

### 4.3. Surface charge distribution analysis

The PyMOL Plugin APBS (Adaptive Poisson-Boltzmann solver) electrostatics software to visualize the distribution of acidic (negatively charged) and basic (positively charged) amino acids on the surface of lumican and TLR9 3D structures. The preparation of molecules including adding missing molecules and assigning atomic charges was automated by pdb2pqr, and grid spacing for APBS calculation was 0.50, which are the default settings. The following parameters were used to run APBS electrostatics to calculate surface charge of lumican and TLR9:

- Prepare molecule with method: pdb2pqr
- Calculate map with APBS with Grid spacing: 0.50
- Molecular surface visualization with electrostatic potential range (+/-): 5.00

### 4.4. Molecular modeling of CpG ODN_2395

The model structure of Class C CpG ODN_2395 was built using BIOVIA Discovery Studio (version 4.5) biopolymer tools. Given that TLR9 demonstrated strong binding and immunostimulatory activity to single stranded CpG DNA, we built a single stranded CpG ODN_2395 as a B-form using standard helix settings.

### 4.5. Docking prediction

We used the HDOCK server for protein-ssDNA interaction. HDOCK server searches for sequence similarity for query receptor and ligand against PDB database for homology modeling and homologous template searching, followed by the prediction of putative binding sites using HDOCKIite global docking algorithm that uses the homologous template for accurate docking prediction (41). The best homologous complex template is chosen based on the highest sequence coverage, highest sequence similarity, and highest resolution available for the docking algorithm. Since the 3D structure of TLR9 ECD, lumican and the ligand CpG ODN_2395 were available, we used the PDB structure as an input. Although TLR9 in complex with CpG ODN_1668 12mer exists in PDB database, the nucleotide sequence coverage was 35%, which only 8 nucleotide sequences were aligned out of 22 of CpG ODN_2395, and not sufficient for the template-based docking prediction. The docking prediction was conducted with the following settings:

- ligand type: ssDNA
- receptor/ligand format: PDB file.
- Docking type: global docking (default)

The interface residues were determined within the distance 5.0A between the receptor and the ligand. The prediction generated 100 complexes. In this study, only topi with the highest confidence and lowest docking score was considered for the analysis.

### 4.6. Protein-DNA interface analysis

The Protein-DNA complex and residues at the Interface were obtained from HDOCK server and visualized in PyMOL. The interface area of the protein-CpG DNA complex was determined by buried surface area (BSA), which is the sum of the accessible surface area (ASA) of unliganded protein and free CpG DNA and subtracted from that of the protein-CpG DNA complex. The buried surface area of each individual residue was calculated from the ASA of unbound state of the individual residue subtracted from the ASA of CpG ODN-bound state of the individual residue. The buried surface area of the protein, CpG DNA, and protein-CpG DNA complex was calculated using PyMOL built-in command line Get_area. The probe diameter was set to 1.4 A (26), which is the diameter of water molecule.

The following equation was used to evaluate the BSA and loss of accessible surface area of individual residue:

- Total Buried Surface Area(A^2^) = (Accessible Surface Area_pro_tein + Accessible Surface Areac_P_G_DNA) - (Accessible Surface Area_P_rotein-cpG dna)
- Buried Surface Area of individual residue (%) = (accessible surface areaFree_residue - accessible surface areaBound_residue)/accessible surface areaFree_residue

The hydrogen bonding, salt-bridges, and hydrophobic interactions in protein-DNA interface were analyzed using Protein-Ligand Interaction Profiler (PLIP) (42). PLIP uses four steps of algorithm to detect interaction patterns: structure preparation to add missing atoms, functional characterization to detect hydrogen bonding acceptor/donor atoms and hydrophobic molecules, rule-based matching based on geometric criteria, and filtering of redundant interactions. The complex generated from the HDOCK server was submitted to PLIP as a PDB format and the interaction data were visualized in PyMOL.

## Supplementary Materials

The following supporting information are included with this manuscript: Figure S1: Pair-wise sequence alignment between lumican and TLR9; Figure S2: Schematic model showing the classes of LRR motifs in Lumican and TLR9; Figure S3: Acidic and basic interface residues in the concave surface of Lumican and TLR9; Figure S4: A ribbon model showing all the TLR9 residues that interacts with the 12 mer CpG ODN_1668; Figure S5: CpG DNA interacting residues in Lumican and TLR9 with similar biochemical property as predicted by Pairwise Sequence Alignment score.

## Author Contributions

Conceptualization: T.C., G.M. and S.C.; Funding acquisition: S.C.; Supervision: G.M. and S.C.; Investigation: T.C. and G.M.; Methodology & Software: T.C.; Analysis: T.C., G.M. and S.C.; Writing-original draft, review and editing: T.C., G.M. and S.C. All authors have read and agreed to the published version of manuscript.

## Institutional Review Board Statement

Not applicable

## Informed Consent Statement

Not applicable

## Data availability Statement

The data and materials used to support the findings of this study are available from the corresponding authors (George Maiti or Shukti Chakravarti) upon request.

## Supplementary Information for

This supplementary document includes the following figures: Figure.S1 to S5

**Figure S1:**
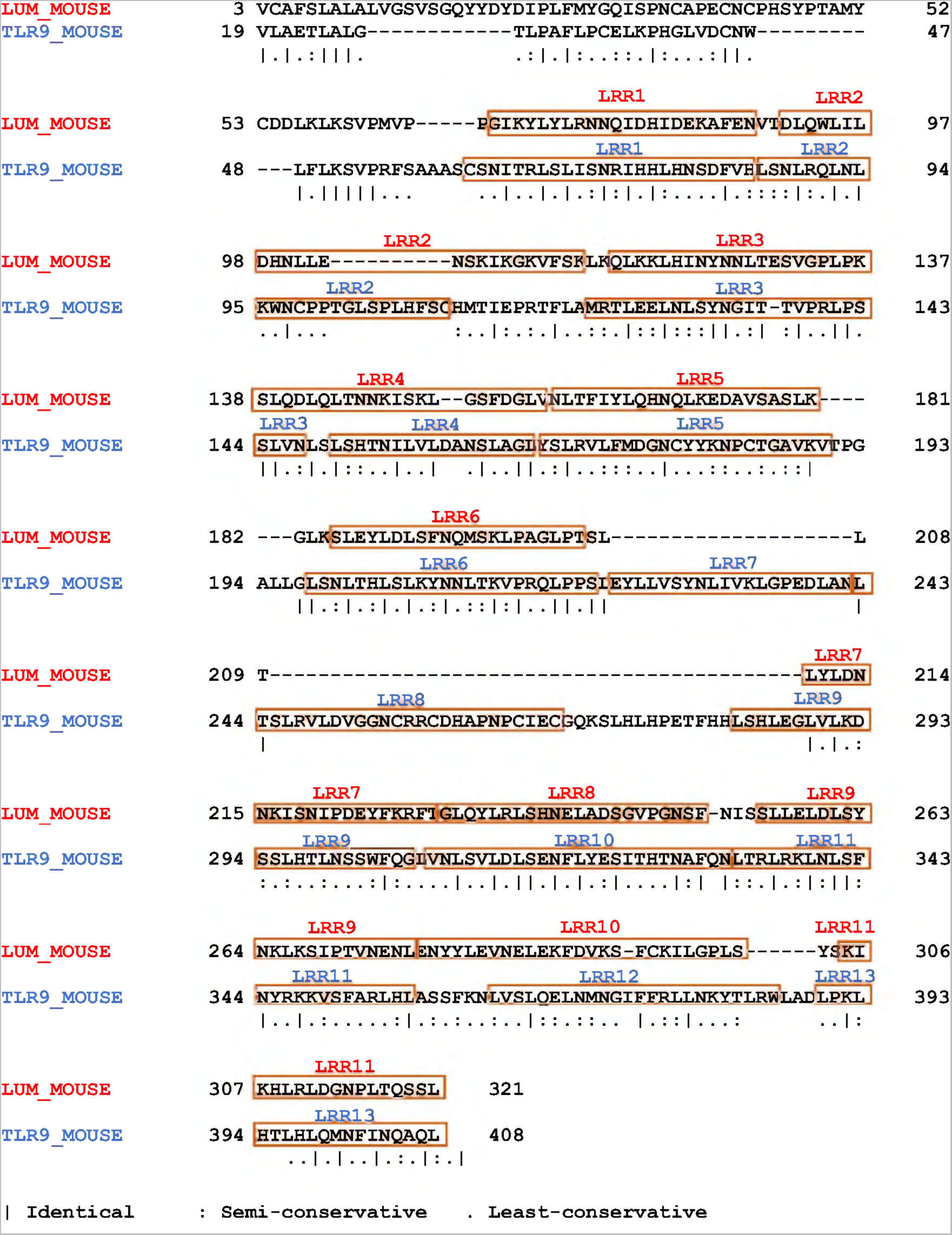
Pair-wise sequence alignment between lumican and TLR9 ECD. The Clustal Omega pairwise sequence alignment between lumican and TLR9 ECD indicates 39.8% and 22.2% sequence similarity and identity, respectively.

**Figure S2:**
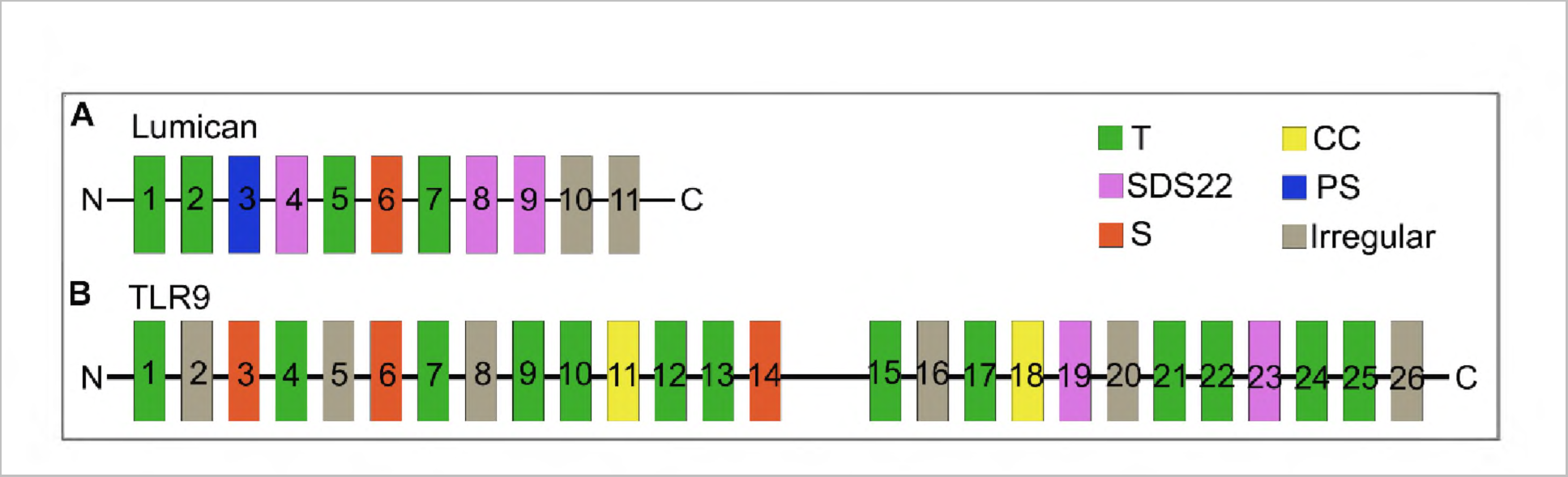
Schematic model showing the classes of LRR motifs in Lumican and TLR9. Both lumican and TLR9 contain a mix of T, S, SDS22 and PS classes of LRRs. **A-B.** The LRR1-11 in lumican mostly consist of T and SDS22 class **(A)** whereas in TLR9 it is mostly T and irregular **(B)**.

**Figure S3:**
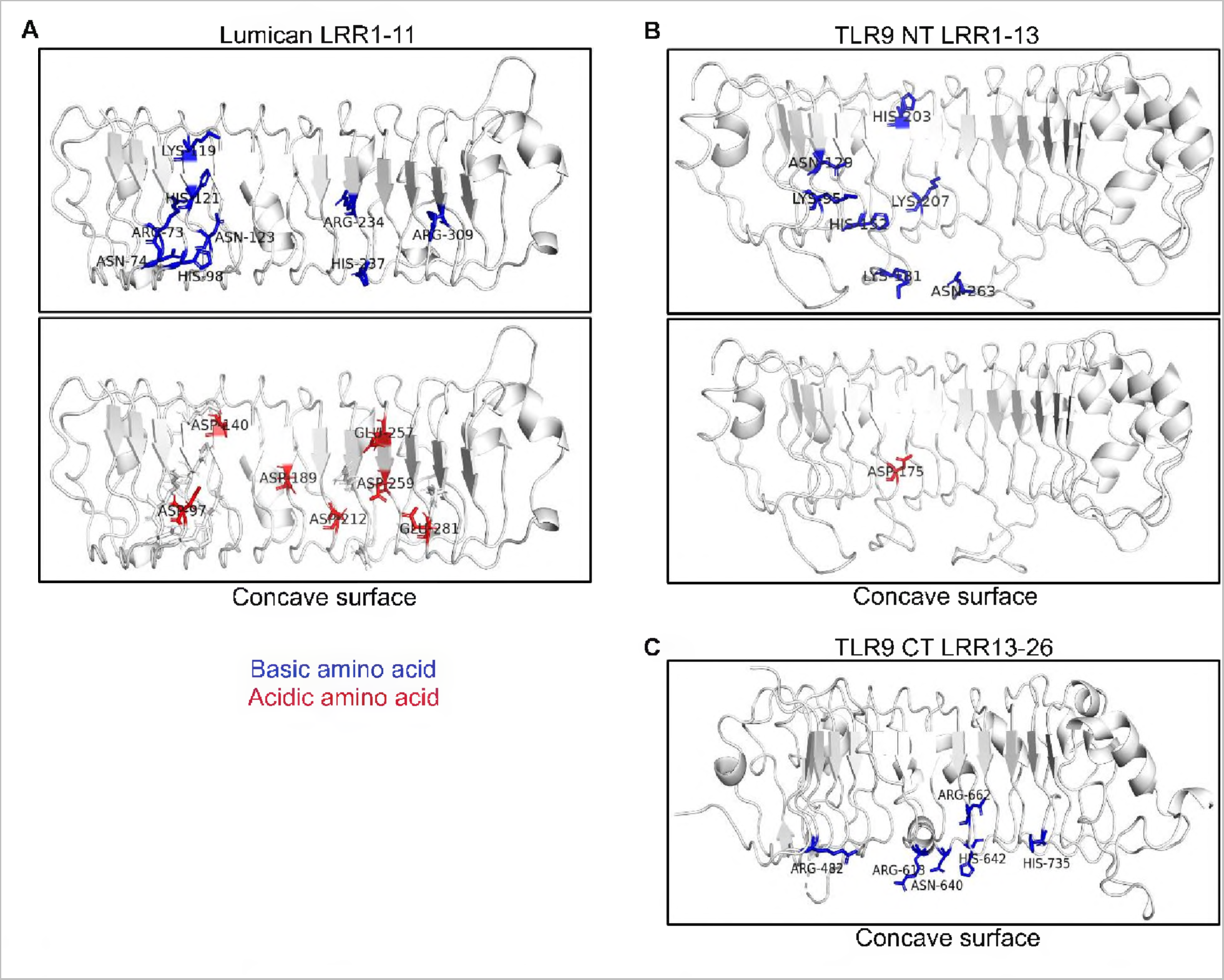
Acidic and basic interface residues in the concave surface of Lumican and TLR9. **A-C.** The electrostatic view shows the acidic residues (red) and basic residues (blue) within the concave surface of lumican **(A)**, TLR9 NT **(B)** and TLR9 CT **(C)**,that interact with the CpG ODN_2395.

**Figure S4:**
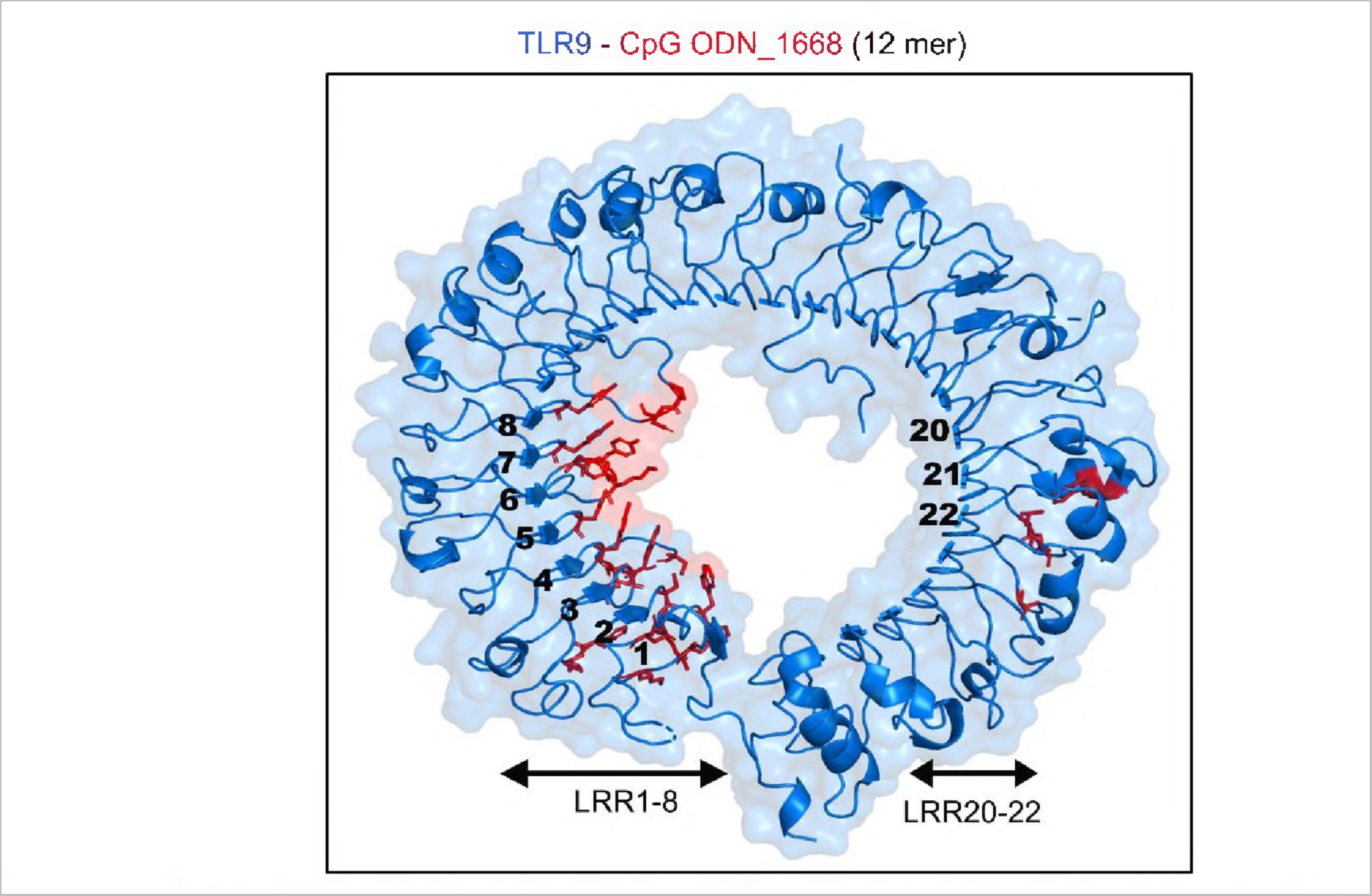
Ribbon model showing all the TLR9 residues that interacts with the 12 mer CpG ODN 1668. The model shows all the interface residues (red) in TLR9 LRR 1-8 and LRR 20-22 that interact with the shorter 12 base CpG ODN_1668.

**Figure S5:**
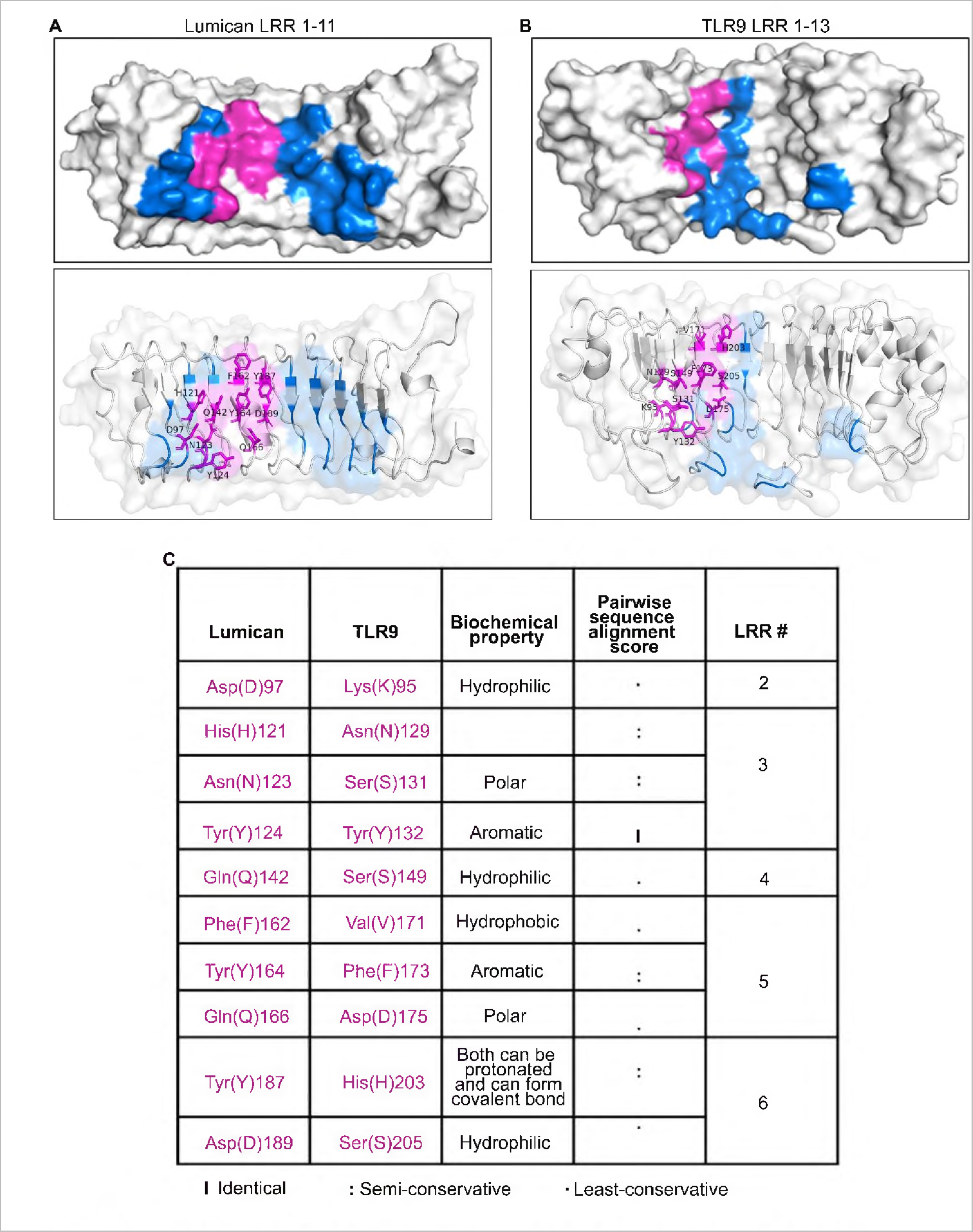
CpG DNA interacting residues in Lumican and TLR9 along with their Pairwise Sequence Alignment score. **A-B.** Surface view showing the CpG ODN_2395 interacting residues (blue and pink) in the concave surface of lumican LRR1-11 **(A)** and TLR9 LRR1-13 **(B)**. **C.** Table shows the list of residues (pink) with their pairwise sequence alignment scores from lumican and TLR9 Clustal Omega alignment.

